# Interphase Human Chromosome Exhibits Out of Equilibrium Glassy Dynamics

**DOI:** 10.1101/193375

**Authors:** Guang Shi, Lei Liu, Changbong Hyeon, D. Thirumalai

**Affiliations:** Biophysics Program, Institute for Physical Science and Technology, University of Maryland, College Park, MD 20742; Korea Institute for Advanced Study, Seoul 02455, Republic of Korea; Department of Chemistry, University of Texas at Austin, Texas, 78712; Corresponding author

## Abstract

The structural organization of the condensed chromosomes is being revealed using chromosome conformation capture experiments and super-resolution imaging techniques. Fingerprints of their three-dimensional organization on length scale from about hundred kilo base pairs to millions of base pairs have emerged using advances in Hi-C and super-resolution microscopy. To determine the poorly understood dynamics of human interphase chromosomes, we created the Chromosome Copolymer Model (CCM) by representing the chromosomes as a self-avoiding polymer with two loci types corresponding to euchromatin and heterochromatin. Using advanced clustering algorithms we establish quantitatively that the simulated contact maps for chromosomes 5 and 10 and those inferred from Hi-C experiments are in agreement. Ward Linkage Matrix (WLM), constructed from spatial distance information, shows that the Topologically Associated Domains (TADs) and compartments predicted from simulations are in agreement with inferred WLM computed using data from super-resolution microscopy experiments. Glassy dynamics is manifested in the stretched exponential relaxation of the structure factor and caging in the mean square displacement of individual loci, ∆_*i*_(*t*) ∼ *t*^*α*^ with 0 < *α* < 1. Remarkably, the distribution of *α*, is extremely broad suggestive of highly heterogeneous dynamics, which is also reflected in the large cell-to-cell variations in the contact maps. Chromosome organization is hierarchical involving the formation of chromosome droplets (CDs) on short genomic scale followed by coalescence of the CDs, reminiscent of Ostwald ripening. We propose that glassy landscapes for the condensed active chromosomes might provide a balance between genomic conformational stability and biological functions.

## Introduction

The organization of chromosomes without topological entanglement or knot formation in the crowded tight space of the nucleus is remarkable. Understanding the structural organization and the dynamics of eukaryotic chromosomes and the mechanism of chromosome territories formation may hold the key to enunciating genome functions (1, 2). Glimpses into the structures of the chromosomes have emerged, thanks to spectacular advances in Chromosome Conformation Capture (3C, 4C, 5C, and Hi-C) experiments (3–5), from which the probability, *P* (*s*), that two loci separated by a certain genomic distance (*s*) are in contact can be inferred. The set of *P* (*s*) as a function of *s*, which is a very low dimensional representation of the spatial organization, constitutes the contact map. More recently, imaging methods like (6) as well as super-resolution technique (7, 8) have more directly determined the positions of the loci of single chromosomes, thus providing a much-needed link to the indirect measure of spatial organization revealed through contact maps. The experiments by Zhuang and coworkers and others (6–8) are of great value because the number of constraints needed to unambiguously infer structures from contact maps alone is very large (9).

Contact maps, constructed from Hi-C experiments, revealed that chromosome is organized into compartments on genomic length scales exceeding megabases (Mbps) (4, 5). The partitioning of the structure into compartments are highly correlated with histone markers of the chromatin loci (5), implying that contacts are enriched within a compartment and depleted between different compartments. The loci associated with active histone markers and those associated with repressive histone markers localize spatially in different compartments. Higher resolution Hi-C experiments (5) have also identified Topologically Associated Domains (TADs) on scales on the order of hundreds of kilobases (3). The TADs are the square patterns along the diagonal of the contact maps in which the probability of two loci being in contact is more probable than between two loci belonging to distinct TADs. super-resolution imaging experiments (6) show most directly that TADs belonging to distinct compartments are spatially separated in the chromosomes.

The experimental studies have inspired a variety of polymer models (10–25), which have provided insights into many aspects of chromosome organization. These studies are particularly important because finding a unique solution (if one exists) to the inverse problem of deriving spatial structures from contact maps is most challenging (9). Some of the features in the contact maps, such as the contact probability, *P* (*s*) may be computed using a homopolymer model, without accounting for epigenetic states, whereas fine structures such as TADs and compartments require copolymer or heteropolymer models (18, 20, 21).

Biological functions, such as the search for genes by transcription factors or mechanism for DNA damage repair, not only depend on genome structure but also the associated dynamics. The use of polymer models, in describing chromatin structure has a rich history (10, 11). More recent studies show that polymer physics concepts have been most useful in predicting the probabilistic nature of chromosome organization inferred from Hi-C experiments (12–23). In contrast, the dynamic aspects of the interphase chromosome have received much less attention (24–29). Experiments have revealed that genome-wide chromatin dynamics (29–32) of chromatin fiber in mammalian cells exhibit heterogeneous sub-diffusive behavior. Thus, it is important to understand how the slow dynamics of the individual locus and long length scale coherent collective motions emerge from the highly organized chromosomes.

Here, we develop a copolymer model to describe both the structure and dynamics of human interphase chromosomes based on the assumption that the large-scale organization of human interphase chromosome is largely driven and maintained by the interactions between the loci of similar epigenetic states. Similar models, that differ greatly in details, have been developed to model the 3D structure of Drosophila chromosomes (18, 20). Jost et al (18) used a heteropolymer model to describe the formation of TADs in Drosophila genome where four different types of monomers representing active, Polycomb, HP-1 and black chromatin, respectively. In a similar spirit, Michieletto et. al, (27) constructed a heteropolymer with three epigenetic states (active, inactive, and acetylated) to probe how the epigenetic states are maintained. A very different reverse-engineering approach, with Hi-C contact maps as inputs, was used to construct an energy function with twenty-seven parameters (21). We took a “bottom-up” approach to incorporate the epigenetic states into the polymer model similar in spirit to the previous studies (18, 27). We show that in order to capture the structural features faithfully, at least two types of beads, representing active and repressive loci are needed. Simulations of the resulting Chromosome Copolymer Model (CCM) for human interphase chromosomes 5 and 10 show that the major structural characteristics, such as the scaling of *P* (*s*) as a function of *s*, compartments, and TADs indicated in the Hi-C contact maps are faithfully reproduced. We use sophisticated clustering algorithms to quantitatively compare the simulated contact maps and those inferred from Hi-C experiments. The compartment feature noted in the Hi-C contact map is due to micro-phase separation between chromosome loci associated with different epigenetic states, implying that a copolymer model is needed for characterizing large-scale genome organization. The TADs emerge by incorporating experimentally inferred positions of the loop anchors, whose formation is facilitated by CTCF motifs. The only free parameter in the CCM, which is the optimal loci-loci interaction strength between loci belonging to distinct epigenetic states, is adjusted to give a good description of the Hi-C contact map. Using simulations based on the resulting CCM we show that chromosome dynamics is highly heterogeneous and exhibits many of the characteristics of out of equilibrium glassy dynamics including stretched exponential decay of the scattering function (*F*_*s*_(*k, t*)), a non-monotonicity behavior in the time dependence of the fourth order susceptivity associated with fluctuations in *F*_*s*_(*k, t*). Of particular note is the remarkable cell-to-cell and loci-to-loci variation in the time (*t*) dependence of the mean square displacement, ∆_*i*_(*t*), of the individual loci. The distribution, *P* (*α*) of the exponent associated with the increase in ∆_*i*_(*t*) ∼ *t*^*α*^ is broad with the simulated and measured *P* (*α*) being in excellent agreement. Our work shows that chromosomes structures are highly dynamic exhibiting large cell-to-cell variations in the contact maps and dynamics. The rugged chromosome energy landscape, with multiple minima separated by large barriers, is perhaps needed to achieve a balance between genomic conformational stability and dynamics for the execution of a variety of biological functions.

## Results

### Choosing the energy scale in the Chromosome Copolymer Model (CCM)

We fixed *N*, the size of the copolymer to *N* = 10, 000, modeling a 12 Mbps (megabases) chromatin fiber, corresponding to a selected region of the Human Cell line GM12878 Chromosome 5 (Chr 5) from 145.87 Mbps to 157.87 Mbps. In the CCM (Fig.1A and Fig.S1), the only unknown parameter is *ϵ*, characterizing the strength of the interaction between the loci (Table I in the SI). We chose a *ϵ* value that reproduces the contact maps that is near quantitative agreement with the Hi-C data. As *ϵ* increases the structures of the chromosome are arranged in such a way that segments with small genomic distance *s* are more likely to be in spatial proximity (see the section **Chromosome Structures in terms of Ward Linkage Matrix (WLM)** below). This is illustrated in Fig.S4, which shows that higher values of *ϵ* lead to clearer segregation between the loci with different colors. The colors encode the genomic locations. The snapshots of the organized chromosome, the good agreement between the calculated and Hi-C contact maps, and the accurate description of the spatial organization as assessed by the Ward Linkage Matrix (WLM) (see the section **Chromosome Structures in terms of Ward Linkage Matrix (WLM)** below and the SI) confirm that *ϵ* = 2.4*k*_B_*T* produces the closest agreement with experiments. Increasing *ϵ* beyond 2.4*k*_B_*T* leads to a worse description of segregation between loci with distinct epigenetic states.

**FIG. 1:**
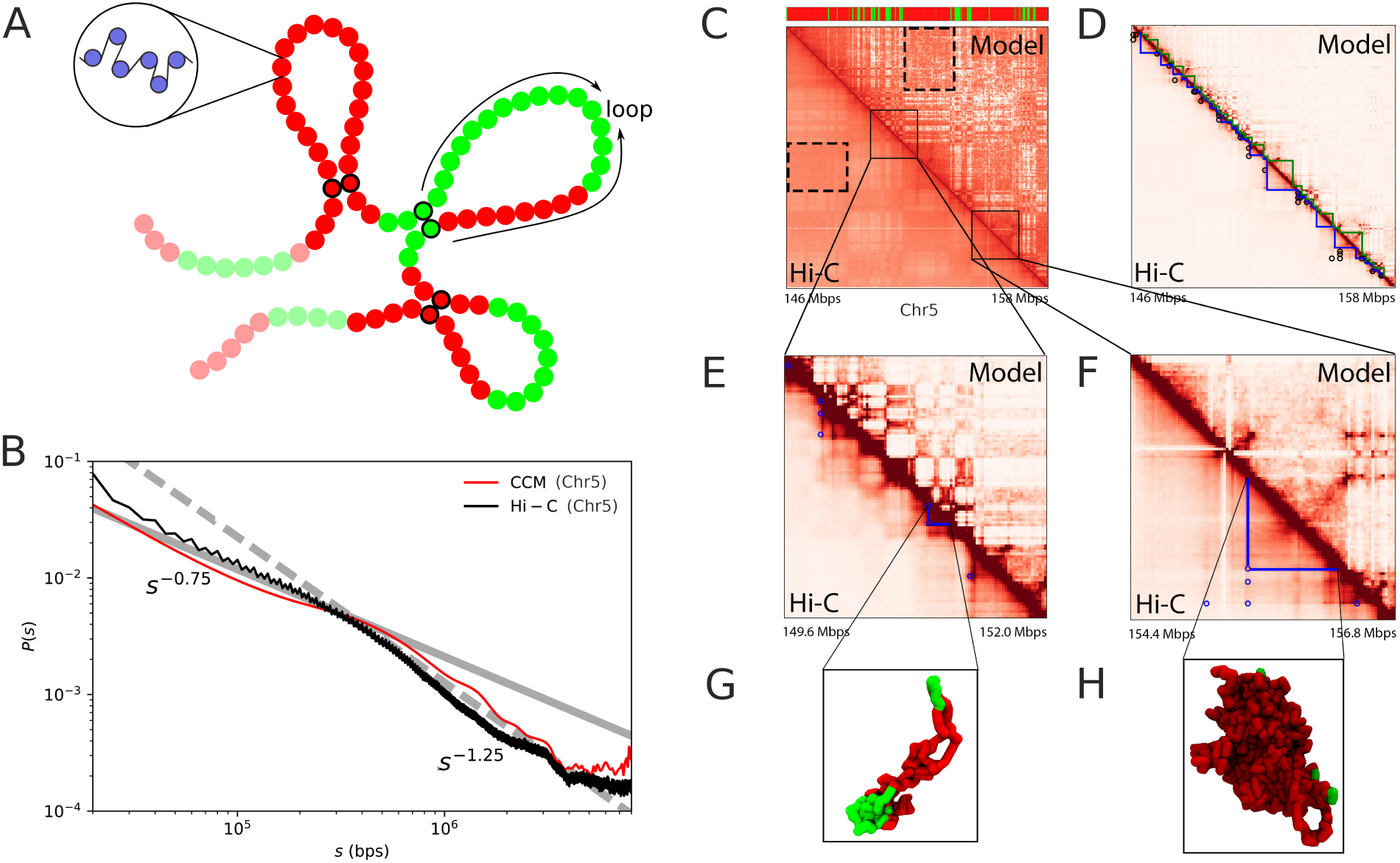
(A) A sketch of the Chromosome Copolymer Model (CCM) Each bead represents 1,200 basepairs (representing roughly six nucleosomes connected by linker DNAs). Red (Green) corresponds to repressive (active) chromatin. The three pairs of loop anchors (in this cartoon) are marked by beads with black boundaries. **(B)** Comparison between experimental data (5) (black) and simulated *P* (*s*). Dashed and solid lines are plots of *s*^*−*1.25^ and *s*^*−*0.75^, respectively. The crossover point between the two scaling regimes at *s^∗^ ∼* 3 *•* 10^5^bps is noticeable in both the experimental and simulated results. **(C)** Comparison of the contact maps inferred from Hi-C experiment (5) (lower triangle) and obtained from simulations (upper triangle) results. For easier visualization, the values of the contact probability are converted to a log_2_ scale. The bar above the map marks the epigenetic states with green (red) representing active (repressive) loci. The dashed black box is an example of a compartment. Such compartment-like structures emerge due to contacts between loci separated by large genomic distances giving rise to spatial order in the organized chromosome. **(D)** Illustration of Topologically Associated Domains (TADs). The blue and green triangles are from experiments and simulations, respectively. The black circles mark the positions of loops detected from experiment data, which are formed by two CTCF motifs. **(E)** The zoom in of the diagonal region for the chromosome segment between 149.6 Mbps to 152.0 Mbps. The blue circle marks the positions of CTCF loops found in the experiment (5). **(F)** Same as **(E)** except for 154.4 Mbps to 156.8 Mbps. **(G)** and **(H)**. Snapshots of two TADs, marked by the blue triangles in **(E)** and **(F)**, respectively. Green (Red) represent euchromatin (heterochromatin).

Furthermore, *P* (*s*) as a function of *s* obtained in simulations with *ϵ* = 2.4*k*_B_*T* is also consistent with experiments (see below). The *s*-dependent contact probability, *P* (*s*) in Fig.1B, shows that there are two scaling regimes. As *ϵ* increases, the probability of short-range (small *s*) increases by several folds, while *P* (*s*) for large *s* decreases by approximately an order of magnitude (Fig.S13A). In particular, for *ϵ* = 1.0*k*_B_*T*, *P* (*s*), decreases much faster compared to experiments at small *s*. In contrast, we find that at *ϵ* = 2.4*k*_B_*T*, *P*(*s*) ∼ *s*^−0.75^ for *s* < 0.5 Mbps and when *s* exceeds ∼ 0.5 Mbps, *P* (*s*) ∼ *s*^−1.25^ (red curve in Fig.1B). Such a behavior, with *P* (*s*) exhibiting two distinct scaling regimes, is in good agreement with experiments (black line in Fig.1B). It is worth pointing out that the two-scaling regimes in *P* (*s*) is a robust feature of all 23 Human interphase chromosomes (Fig.S14).

### Active and repressive loci micro-phase segregate

Comparison of the contact maps between simulations and experiments illustrates that compartment formation appearing as plaid or checkerboard patterns in Fig.1C, demonstrating good agreement with Hi-C data (4, 5). The dashed rectangles mark the border of one such compartment enriched predominantly with interactions between loci of the same type, suggesting that compartments are formed through the clustering of the chromatin segments with the same epigenetic states. Interestingly, a previous experimental study suggests that the chromatin structuring in Topologically Associated Domains (TADs) is also driven by the epigenome feature (33). In order to make the comparison precise, we treated the contact maps as probabilistic matrices and used a variety of mathematical methods to quantitatively compare large matrices. First, the checkerboard pattern in the contact map is more prominent when illustrated using the Spearman correlation map (see SI for details and Figs.S6 and S7). Second, to quantitatively compare the simulated results with experiments, we use the spectral co-clustering algorithm (34) to bi-cluster the computed Spearman correlation map (see SI for details). Finally, the similarity between the simulated and experimental data is assessed using the Adjusted Mutual Information Score (AMI), which is also described in the SI. The CCM model, based only on epigenetic information and the locations of the loop anchors, yields an AMI score that results in correctly reproducing *≈* 81% of the compartments obtained from the experimental data. In contrast, a pseudo homopolymer model with *ϵ*_*AA*_ = *ϵ*_*BB*_ = *ϵ*_*AB*_ = *ϵ*), which preserves the nature of epigenetic states, (see SI), has an absolute AMI score that is 200 times smaller (Fig.S9), does not lead to the formation of compartments (correctly reproducing only *≈* 51% of the compartments, no better than random assignments). Thus, the CCM is the minimal model needed to reproduce the essential features found in the contact map.

The inset in Fig.2A, displaying a typical snapshot of the condensed chromosome, reveals that active (A, green) and repressive (B, red) loci are clustered together, undergoing micro-phase separation (see Methods for definition of active and repressive loci). The tendency to segregate is vividly illustrated in the radial distribution functions *g_AA_* (*r*), *g_BB_* (*r*) and *g_AB_* (*r*), which shows (Fig.2A) that *g_AA_* (*r*) and *g_BB_* (*r*) have much higher values than *g_AB_* (*r*) implying that active and repressive loci form the clusters of their own, and do not mix. Such a micro-phase separation between the A-rich and B-rich regions directly gives rise to compartments in the contact map. Interestingly, the normalized radial density (Fig.2B) shows that active chromatin exhibits a peak at large radial distance, *r* implying that the active loci localize on the periphery of the condensed chromosome whereas repressive chromatin is more homogeneously distributed. Visual inspection of the simulation trajectories also suggests that active and repressive chromatins are often separated in a polarized fashion, in accord with a recent experimental study (6), which shows that the two compartments are indeed similarly spatially arranged.

**FIG. 2:**
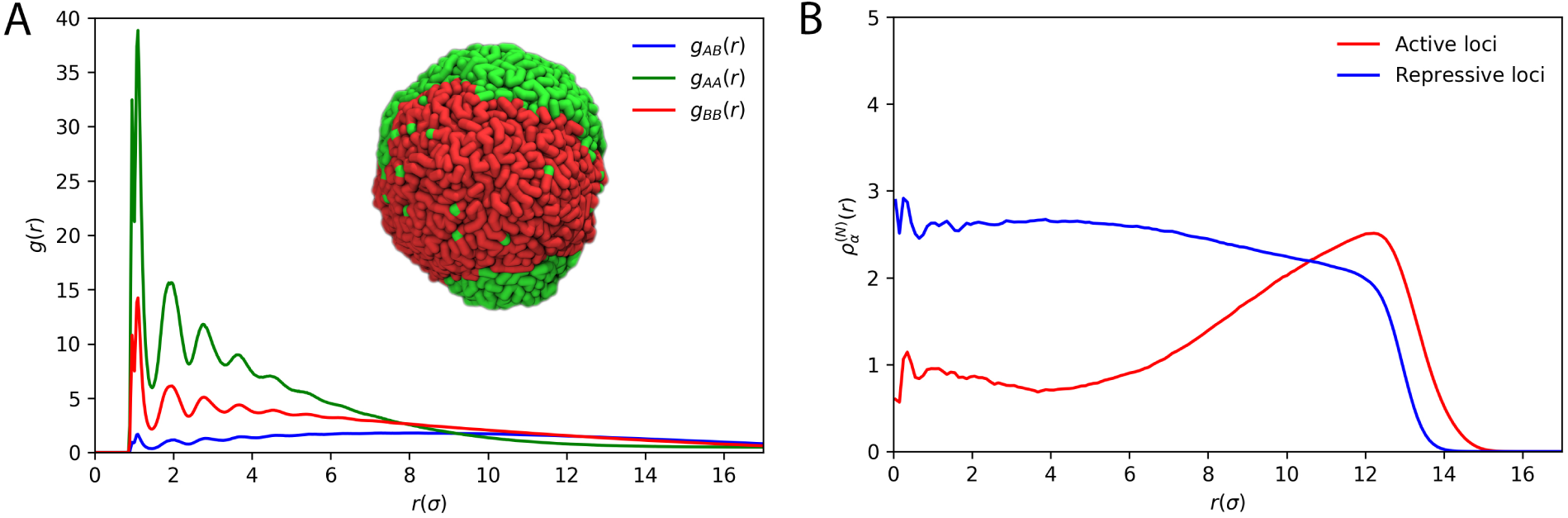
**(A)** Radial distribution functions, *g*(*r*), as a function of *r* (in the unit of *σ*) between active and repressive loci. The inset shows the typical conformation of the compact chromosome. Green and red segments correspond to active and repressive loci, respectively. The structure vividly reveals micro-phase separation between active and repressive loci. **(B)** The normalized radial density, 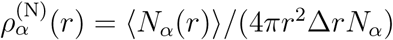, where *N*_*α*_ (*r*) is the number of loci of given type *α*found in the spherical shell between *r* and *r*+∆*r*, *N*_*α*_ is the total number of loci of that type. The bracket *•O* is the ensemble average, *V* is the volume of the globule, given by 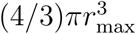 where *r*max = 17*σ*; 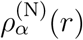 shows that the active loci are predominantly localized on the periphery of the condensed chromosome. The repressive loci are more uniformly distributed.

### Spatial organization of the compact chromosome

In order to illustrate the spatial organization of the chromosome, we introduce the distance function,

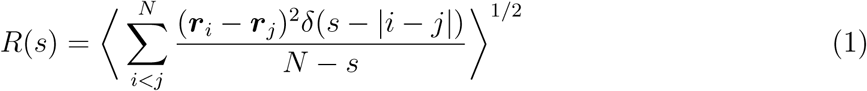

where 〈⋅〉 denotes both an ensemble and time average. We calculated *R*(*s*), the mean end-to-end distance between the loci, by constraining the genomic distance *|i − j|* to *s*. If the structured chromosome is maximally compact on all length scales, we expect *R*(*s*) ∼ *s*^1/3^ for all *s*. However, the plot of *R*(*s*) on a log-log scale shows that in the range 10^5^ ≲ *s* ≳ 10^6^ bps, *R*(*s*) ∼ *s*^0.2^. The plateau at large *s* arises due to *s* reaching the boundary of the compact structure. The inset in Fig.3A, comparing the simulation result and experimental data (6), both show the same scaling for *R*(*s*) as a function of *s*.

By a systematic analysis of the FISH data, Wang et al (6) established that the probability of contact formation, *P* (*s*), is inversely proportional to a power of *R*(*s*), with the latter providing a direct picture of the spatial organization. Similarly, in this work, we explored the relation between between *C_ij_* and *R_ij_* where *C_ij_*(= *P_ij_ ∑_i<j_ C_ij_ ∝ P*_*ij*_) is the number of contacts between loci *i* and *j*, and *R*_*ij*_ = 〈(***r**_i_ − **r***_*j*_)^2^〉^1/2^ is the mean distance between them. The heat map of (1*/C_ij_, R*_*ij*_) in Fig.3B shows that the two matrices are proportional to each other. In accord with the FISH data (6), we find that 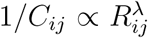 where *λ ≈* 4, suggesting that larger mean spatial distance between loci *i* and *j* implies smaller contact probability, which is the usual, assumption when experimental Hi-C data is used to infer three-dimensional chromosome organization. The decrease of *C*_*ij*_ with increasing *R*_*ij*_ with a large value of *λ*, is unexpected but is an important finding needed to link contact maps and spatial structures.

**FIG. 3:**
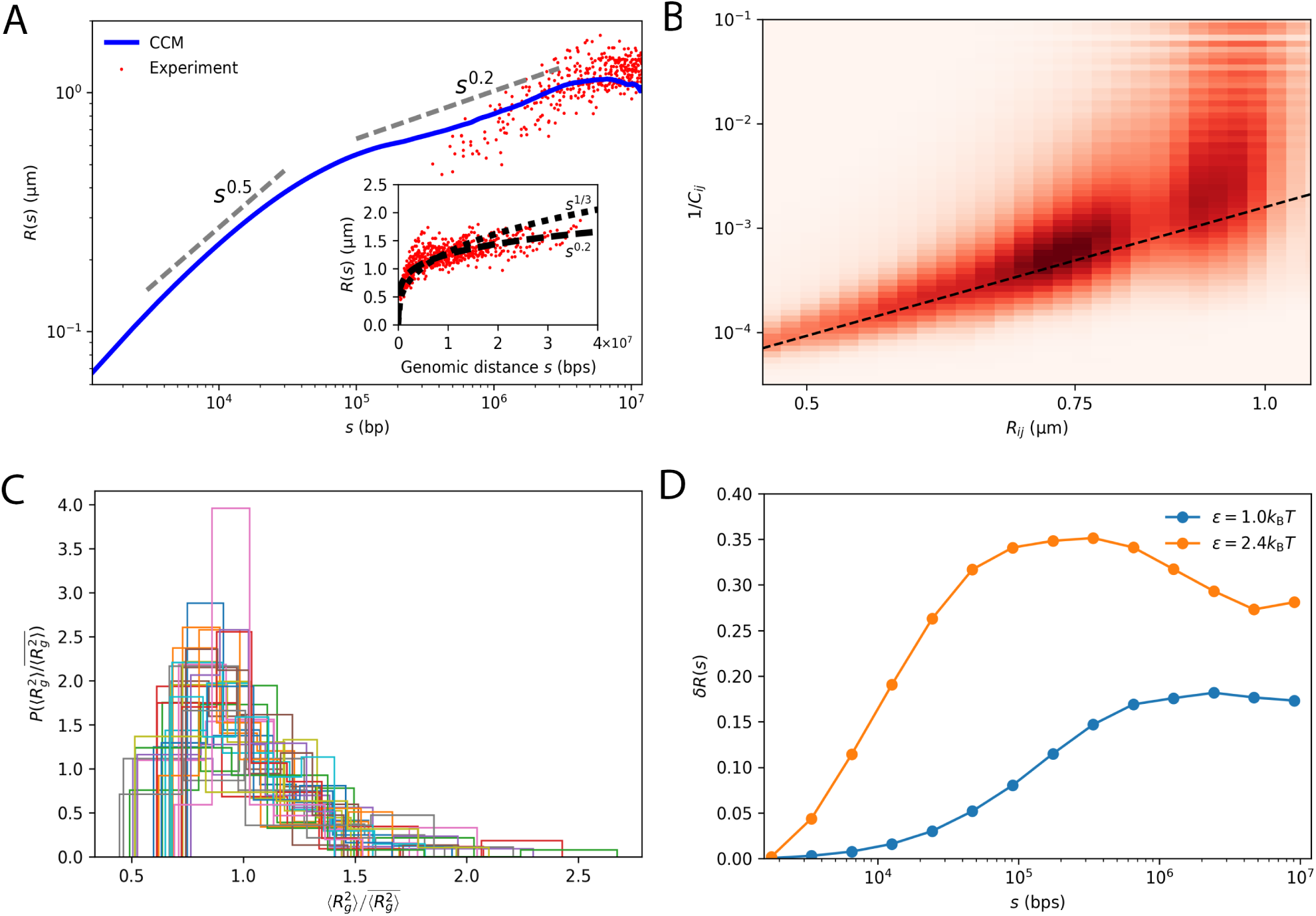
Organization and fluctuations of the chromosome structures. **(A)** The dependence of the spatial distance *R*(*s*) (Eq.1) on the genomic distance, *s*. Grey dashed lines, indicating the slopes, are guides to the eye. The red dots are experimental data taken from Ref. (6) for *s* < 1.2*×*10^7^bps. The inset shows the complete set of experimental data. Short dashed and long dashed lines are *s*^1/3^ and *s*^0.2^, respectively. At small *s* (*s* < 10^5^bps), *R*(*s*) ∼ *s*^0.5^ implying that chromatin behaves as almost an ideal chain. **(B)** The heatmap of the 2D histogram of (*R_ij_,* 1*/C*_*ij*_). The dashed black line is the curve with scaling exponent 4.1, which coincides with the value obtained by fitting the experimental data (6). **(C)** Distribution 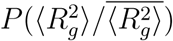, where 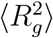 is the time average value of the squared radius of gyration of a single trajectory and 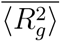is the mean value averaged over all independent trajectories. Different colors represent 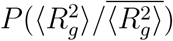 for the thirty-two individual TADs. The distribution is surprisingly wide which suggests that TAD structures vary from cell-to-cell. **(D)** Coefficient of variation *δR*(*s*) = (*R*^2^(*s*)*O − R*(*s*)*O*2)^1/2^*/ R*(*s*)*O*, computed from simulations, shows a non-monotonic dependence on *s* for *ϵ* = 2.4*k*_B_*T*, increasing till *s ∼* 10^5^bps and decreases at larger *s*.

The slope of the dashed line in Fig.3B obtained using the data in Ref. (6), is 4.1, which coincides with our simulation results. Mean field arguments (35) suggest that *P* (*s*) ∼ *R*(*s*)^−3^, which follows from the observation that the end of the chain is uniformly distributed over a volume *R*^3^(*s*). This is neither consistent with our simulations nor with experiments, implying that the distribution of the chain ends is greatly skewed. Although both the simulated and experimental results establish a strong correlation between *R*(*s*) and *P* (*s*), such a correlation is only valid in an ensemble sense (see SI for additional discussions and as well as Ref. (36)).

### Topologically Associated Domains and their shapes

Our model reproduces Topologically Associated Domains (TADs), depicted as triangles in Fig.1D, of an average length of 200 kbps along the diagonal of the contact map in which the interactions between the loci are greatly enhanced. It has been noted (5) that in a majority of cases, boundaries of the TADs are marked by a pair of CTCF motifs with a high probability of interaction between them. They are visualized as peaks in the Hi-C map (Fig.1D). To quantitatively detect the boundaries of the TADs, we adopt the procedure described in Ref. (3) to identify the position of each TAD (see SI for a description of the Directionality Index method for identifying TADs). The boundaries of the TADs, shown in blue (Hi-C data) and green (simulations) are reproduced by the CCM (Fig.1D).

To investigate the sizes and shapes of each individual TADs, we calculated the radii of gyration, *R*_*g*_, the relative shape anisotropies *κ*^2^, as well as the shape parameters, *S*, for 32 TADs (see SI for details). The results are shown in Fig.S10. The mean *R*_*g*_ for each individual TADs scales as their genomic length with exponent 0.27, which is an indicator of the compact structures for the TADs. However, unlike compact globular objects, their shapes are far from being globular and are much more irregular with smaller TADs adopting more irregular shapes compared to the larger TADs (see 〈*κ*^2^〉 and 〈S〉 as a function of TAD size in Fig.S10). Such compact but irregularly shaped nature of TADs are vividly illustrated by typical snapshots for the two TADs (Figs.1G and 1H). How can we understand this non-trivial highly aspherical shapes of the TADs when the chromosome is spherical on long length scales (several Mbps) ? Since TADs are constrained by the CTCF loops, they may be viewed locally as ring polymers. It has been shown (37) that ring polymers in a melt are compact objects but adopt irregular shapes, consistent with our prediction for TADs.

We then wondered if TADs in each individual cells have similar sizes and shapes. We computed the dispersion in *R*_*g*_, *κ* and *S* (Fig.3C and Figs.S10 and S11) among different trajectories. 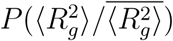, of the mean square radius of gyration 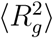 for the thirty-two Chr5 TADs in each trajectory normalized by the average 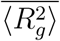 of each individual TAD is shown in Fig.3C. The bracket (bar) is the time (ensemble) average. The large dispersion in 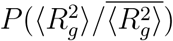 (Fig.3C) as well as 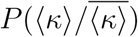 and 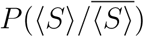 (Fig.S11) suggest that TADs are fluctuating objects, which exhibit substantial cell-to-cell variations. Our result supports the recent FISH (38) and single cell Hi-C experimental findings (39, 40), showing that individual TAD compaction varies widely from highly extended to compact states among different cells. To decipher how the variation of the structure of the chromosome changes as a function of *s*, we calculated the coefficient of variation, 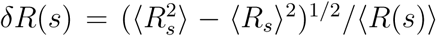. Interestingly, *δR*(*s*) first increases with *s* up to *s ≈* 10^5^ ∼ 10^6^ bps and then decreases as *s* further increases (Fig.3D). Analysis of the experimental data from Ref. (6) shows a similar decreasing trend for *s* > 10^5^bps (Fig.S12(C)). Higher resolution experiments are needed to resolve the variance for *s* < 10^5^bps. The predicted non-monotonic dependence of *δR*(*s*) on *s* is amenable to experimental test.

### Chromosome Structures in terms of Ward Linkage Matrix (WLM)

To quantitatively analyze the spatial organization of the compact chromosome, we use the unsupervised agglomerative clustering algorithm to reveal the hierarchy organization on the different length scales. A different method, which is also based on clustering techniques, has recently been applied to Hi-C contact map (41). We use the Ward Linkage Matrix (WLM), which is directly applicable to the spatial distance matrix, ***R*** in which the element, *R*_*ij*_, is the distance between the loci *i* and *j* (see SI for details). We also constructed the experimental WLM by converting the Hi-C contact map to a distance map by exploiting the approximate relationship between *R*_*ij*_ and 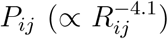 discussed previously (see Fig.3B). The advantages of using distance matrices instead of contact maps are two folds. First, matrix ***R*** is a direct depiction of the three-dimensional organization of the chromosome. Because WLM, constructed from ***R***, is a cophenetic matrix, which can be used to reveal the hierarchical nature of the chromosome organization. Second, the contact map matrix elements do not obey triangle inequality. Therefore, it is not a good indicator of the actual 3D spatial arrangement of the loci. We show the WLM for the two *ϵ* values (Fig.4A) and the comparison between WLM computed based on experimental data and WLM for *ϵ* = 2.4*k*_B_*T* (Fig.4B). Visual inspection of the WLMs for *ϵ* = 2.4*k*_B_*T* shows distinct segregation in the spatial arrangement of the loci. It is clear from Fig.4B that the WLM, reconstructed from Hi-C data, and simulations result with *ϵ* = 2.4*k*_B_*T* are almost identical. From the heat maps of the WLMs show that, for both *ϵ* = 1.0*k*_B_*T* and *ϵ* = 2.0*k*_B_*T* (Fig.S4), we surmise that loci with large genomic separation *s* are in spatial proximity, which is inconsistent with the experimental WLM. The Pearson correlation coefficient between experimental result and CCM using *ϵ* = 2.4*k*_B_*T* is 0.96 (0.53 for *ϵ* = 1.0*k*_B_*T*, 0.84 for *ϵ* = 2.0*k*_B_*T* and 0.75 for *ϵ* = 2.7*k*_B_*T*). Thus, the poorer agreement between the simulated WLM (Fig.S4) as well as Spearman correlation matrix (Fig.S6) using *ϵ* = (1.0, 2.0, 2.7)*k*_B_*T* and experiments, compared to *ϵ* = 2.4*k*_B_*T*, further justifies the latter as the optimum value in the CCM. We find it remarkable that the CCM, with only one adjusted energy scale (*ϵ*, the ratio 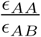) is sufficient to produce such a robust agreement with experiments. We wish to point out that *ϵ* = 2.7*k*_B_*T* does not produce as good an agreement with experiment (see Fig.S4) including the inferior value of the Pearson correlation coefficient mentioned above.

**FIG. 4:**
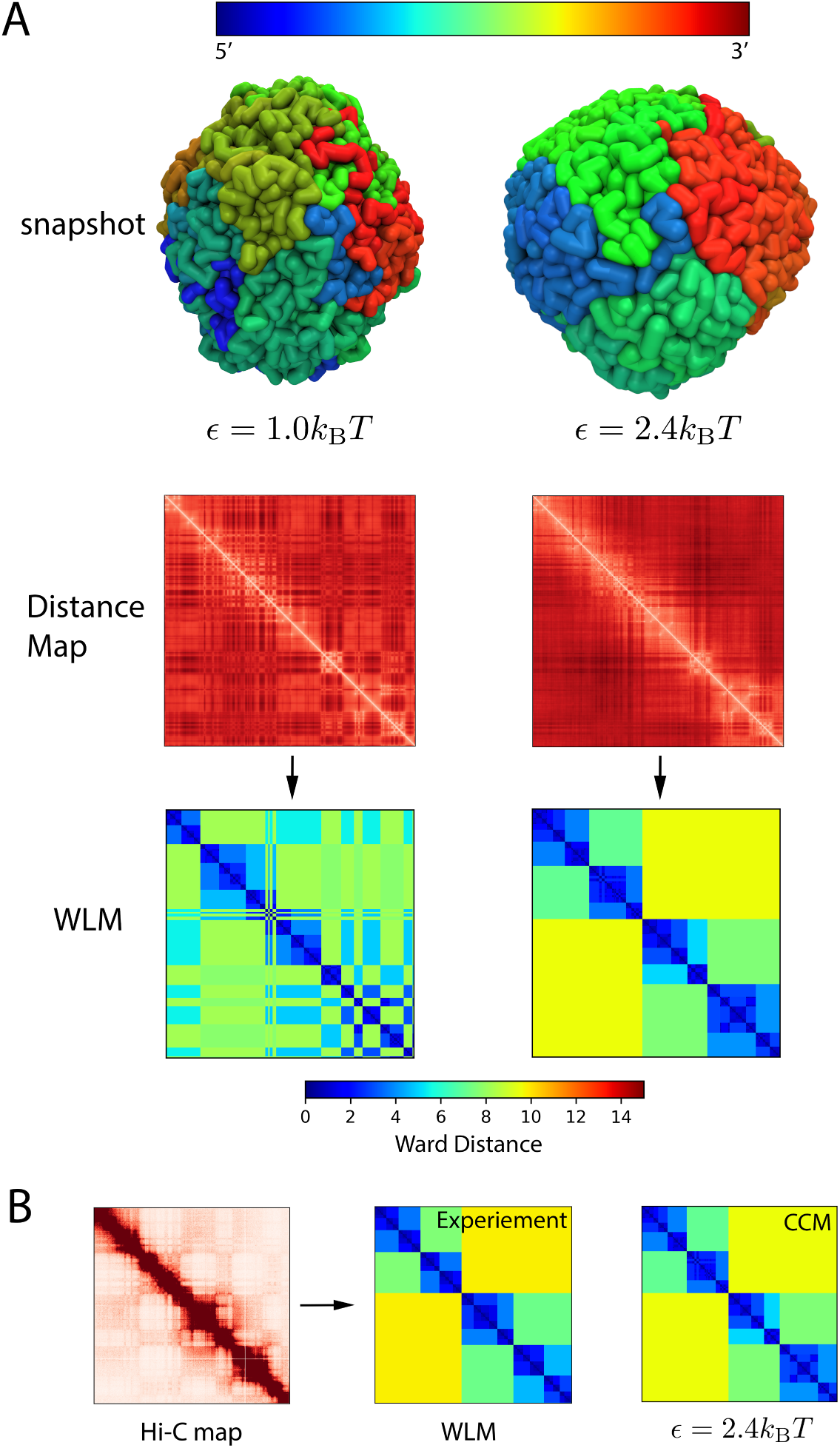
**(A)** Typical conformations of the organized chromosome for *ϵ* = 1.0*k*_B_*T* (left) and 2.4*k*_B_*T* (right). The panel below shows the distance maps (red) and the Ward Linkage Matrix. The color bar at the bottom indicates the value of Ward distance (see SI) **(B)** Comparison of the ensemble average Ward Linkage Matrix (WLM) calculated using the experimental and simulated (*ϵ* = 2.4*k*_B_*T*) contact maps.

### Cell-to-cell variations in the WLM

To assess the large structural variations between cells, we calculated the WLM for individual cells. We obtain the single cell WLM as a time average of a trajectory generated for an individual cell. Fig.5 shows that there are dramatic differences between the WLM for individual cells, with the ensemble average deviating greatly from the patterns in individual cells. Thus, it is likely that the chromosome structure is highly heterogeneous. These findings are reflected in the broad width of the distribution of the Pearson Correlation between all pairs of cells. The mean of the distribution is 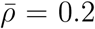 small, implying little overlap in the WLMs between any two cells.

In order to make quantitative comparisons to experimental data, with the goal of elucidating large-scale variations in the spatial organizations of human interphase chromosomes, we constructed single cell WLMs for Chr21 using the spatial distance data provided in (6) and computed the corresponding *P* (*ρ*) (Fig.5B). The results show that the experimentally organization of Chr21 *in vivo* also exhibits large variations manifested by the distribution *P* (*ρ*) covering a narrow range of low values of *ρ* with a small mean 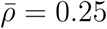. Comparison to simulated result suggest that Chr21 shows a slightly lower degree of structural heterogeneity compared to Chr5 investigated using CCM. Nevertheless, both the simulated and experimental results indicate that human interphase chromosomes do not have any well-defined “native structure”. To investigate whether Chr5 has a small number of distinct spatially structures, we show two-dimensional t-SNE (t-distributed stochastic neighboring embedding) representation of 90 individual WLMs of the metric 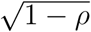 (Fig.5C). It is clear that there is no dominant cluster, indicating that each Chr5 in single cells is organized differently rather than belonging to a small subset of conformational states. Such large cell-to-cell variations in the structures, without a small number of well defined states, is another hallmark of glasses, which are also revealed in recent experiments (40, 42). The presence of multiple organized structures has profound consequences on the chromosome dynamics (see below).

**FIG. 5:**
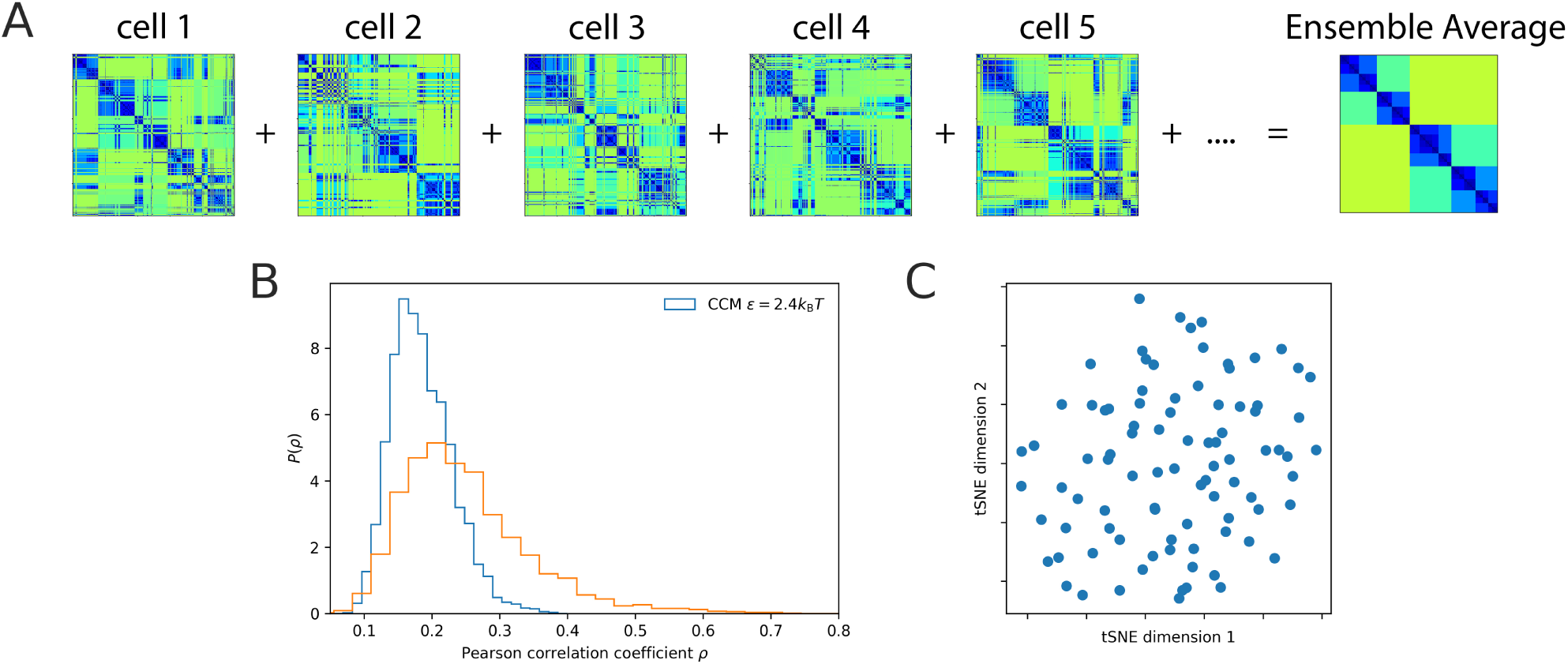
Structural heterogeneity of the compact chromosome. **(A)** Ward Linkage Matrices of different individual cells. The single cell WLM is the time average result over a single trajectory. The ensemble average WLM (rightmost) and the experimental WLM are in clear quantitative agreement (Fig.4B). However, the spatial organization varies from cell to cell. Each cell has very different WLM, implying their structures are distinct. To quantify such heterogeneity, we calculated the Pearson correlation coefficient, *ρ*, between WLMs of any two cells. (**B**) The distribution of *ρ*, *P* (*ρ*), with a mean 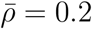 (blue curve). The *P* (*ρ*) distribution, spanning the low range of *ρ* values, is a further demonstration of structural heterogeneity in individual cells. In yellow we plot *P* (*ρ*) with 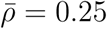 for 120 individual human interphase Chr21, computed using the single cell WLMs constructed from experimental measured spatial distance data provided in (6). **(C)** Two-dimensional t-SNE (t-distributed stochastic neighboring embedding) representation of spatial organization of simulated individual Chr5 on the distance metric 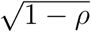.

### Chromosome dynamics is glassy

We probe the dynamics of the organized chromosome with *ϵ* = 2.4*k*_B_*T*, a value that yields the best agreement with the experimental Hi-C contact map. We first calculated the incoherent scattering function, 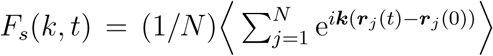 where ***r***_*j*_(*t*) is the position of *j*^*th*^ loci at time *t*. The decay of *F*_*s*_(*k, t*) (orange line in Fig.6A) for *k ∼* 1*/r*_*s*_ (*r*_*s*_ is the position of the first peak in the radial distribution function (*g_AA_* (*r*) and*g_BB_* (*r*)) (Fig.2A)) is best fit using the stretched exponential function, *F*_*s*_(*k, t*) ∼ *e*^−(*t/τα*)*β*^ with a small stretching coefficient, *β ≈* 0.27. The stretched exponential decay with small *β* is one hallmark of glassy dynamics. For comparison, *F*_*s*_(*k, t*) decays exponentially for *ϵ* = 1.0*k*_B_*T*, implying liquid-like dynamics (blue line in Fig.6A).

**FIG. 6:**
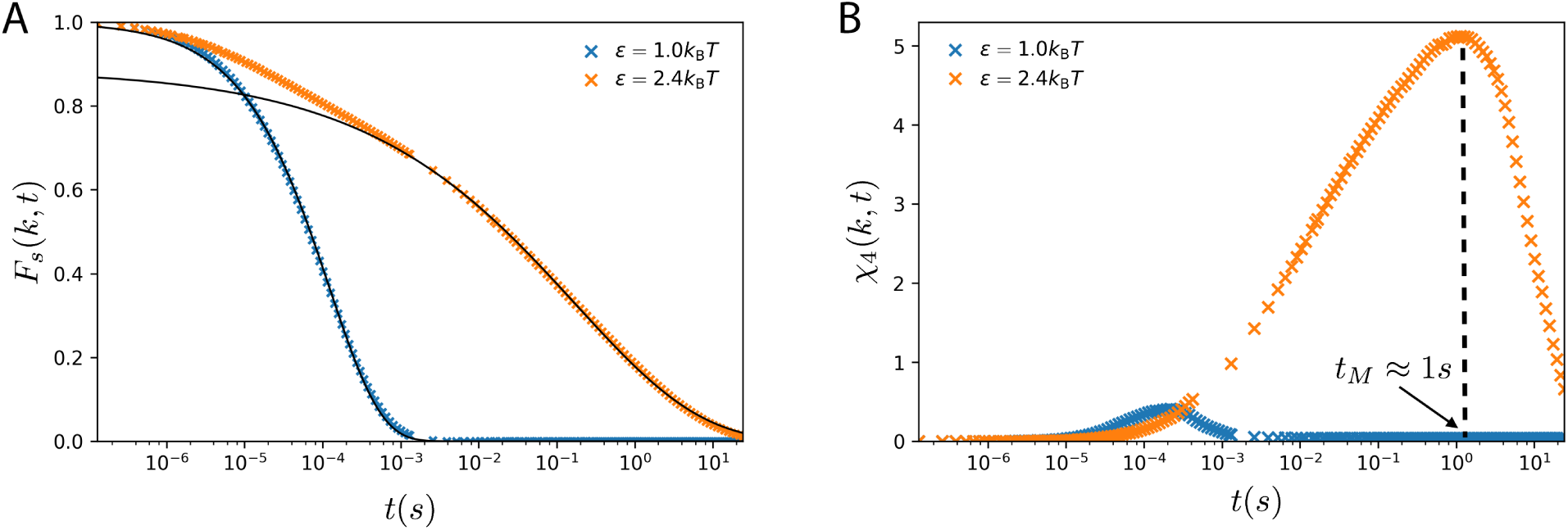
**(A)** Intermediate scattering function obtained for *ϵ* = 1.0*k*_B_*T* (blue) and *ϵ* = 2.4*k*_B_*T* (orange). The line shows an exponential function fit, *F*_*s*_(*k, t*), for *ϵ* = 1.0*k*_B_*T*. For *ϵ* = 2.4*k*_B_*T*, *F*_*s*_(*k, t*) ∼ *ϵ*^−(*t/tα*)*β*^ with *β* = 0.27, for *t* exceeding a few milliseconds (black curve). **(B)** The fourth order susceptibility, *χ*_4_(*t*), used as a function to demonstrate dynamic heterogeneity. The peak in *χ*_4_(*t*) for *ϵ* = 2.4*k*_B_*T* around *t_M_ ≈* 1s is a signature of heterogeneity.

In the context of relaxation in supercooled liquids, it has been shown that the fourth order susceptibility (43), *χ*_4_(*k, t*) = *N*[〈*F*_*s*_(*k, t*)^2^*〉 − 〈F*_*s*_(*k, t*)〉^2^] provides a unique way of distinguishing between fluctuations in the liquid and frozen states. As in structural glasses, the value of *χ*_4_(*k, t*) increases with *t* reaching a peak at *t* = *t*_*M*_ and decays at longer times. The peak in the *χ*_4_(*k, t*) is an indication of dynamic heterogeneity, which in the chromosome is manifested as dramatic variations in the loci dynamics (see below). For *ϵ* = 2.4*k*_B_*T*, *χ*_4_(*k, t*) reaches a maximum at *t_M_ ≈* 1*s* (Fig.6B), which surprisingly, is the same order of magnitude (∼ 5*s*) in which chromatin movement was found to be coherent on a length scale of 1*µ*m (31). The dynamics in *F*_*s*_(*k, t*) and *χ*_4_(*k, t*) together show that Human Interphase chromosome dynamics is glassy (26), and highly heterogeneous.

### Single loci Mean Square Displacements are heterogeneous

In order to ascertain the consequences of glassy dynamics at the microscopic level, we plot the MSD, 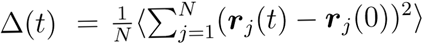 in Fig.7, from which a few conclusions can be drawn.

1. Because of the polymeric nature of the chromosome, the maximum excursion in 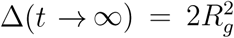, where *R_g_ ≈* 0.7*µ*m is the radius of gyration of Chr 5. Consequently, for both *ϵ* = 1.0*k*_B_*T* (red) and *ϵ* = 2.4*k*_B_*T*, ∆(*t*) in the long time limit is smaller than 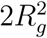. For *ϵ* = 2.4*k*_B_*T* (green), ∆(*t*) shows a crossover at *t ≈* 10^−2^*s* from slow to a faster diffusion, another indicator of glassy dynamics (44). The slow diffusion is due to caging by neighboring loci, which is similar to what is typically observed in glasses. The plateau in ∆(*t*) (Fig.7A) is not pronounced, suggesting that the compact chromosome is likely on the edge of glassiness. The crossover is more prominent in the time-dependence of the mean squared displacement of single loci (see below).
2. The two dashed lines in Fig.7A show ∆(*t*) ∼ *t*^*α*^ with *α* = 0.45. The value of *s* is close to 0.5 for the condensed polymer, which can be understood using the following arguments. The total friction coefficient experienced by the whole chain is the sum of contributions from each of the *N* monomers, 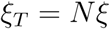. The time for the chain to move a distance *≈ R*_*g*_ is 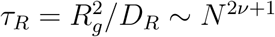. Let us assume that the diffusion of each monomer scales as *Dt*^*α*^. If each monomer moves a distance on the order of *R*_*g*_ then the chain as a whole will diffuse by *R*_*g*_. Thus, by equating 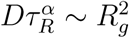, we get *α* = 2*ν/*(2*ν* + 1). For an ideal chain *ν* = 0.5, which recovers the prediction by Rouse model, *α* = 0.5. For a self-avoiding chain, *ν ≈* 0.6, we get *α ≈* 0.54. For a condensed chain, *ν* = 1/3, we get *α* = 0.4, thus rationalizing the findings in the simulations. Similar arguments have been reported recently for dynamics associated with fractal globule (28) and for the *β−*polymer model (29). Surprisingly, *α* = 0.45 found in simulations is in good agreement with recent experimental findings (32). We also obtained a similar result using a different chromosome model (45), when the dynamics were examined on a longer length scale.
3. We also calculated the diffusion of a single locus (sMSD) defined as 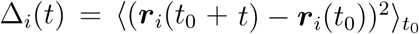, where 〈⋅〉_*t*0_ is the average over the initial time *t*_0_. Distinct differences are found between the polymer exhibiting liquid-like and glassy-like dynamics. The variance in single loci MSD is large for *ϵ* = 2.4*k*_B_*T*, illustrated in Fig.7B, which shows 10 typical trajectories for *ϵ* = 1.0*k*_B_*T* and *ϵ* = 2.4*k*_B_*T* each. For glassy dynamics, we found that the loci exhibiting high and low mobilities coexist in the chromosome, with orders of magnitude difference in the values of the effective diffusion coefficients, obtained by fitting ∆_*i*_(*t*) = *D_α_t^α^_i_*. Caging effects are also evident on the timescale as long as seconds. Some loci are found to exhibit caging-hopping diffusion, which is a hallmark in glass-forming systems (46, 47). Interestingly, such caging-hopping process has been observed in Human cell some time ago (48).
4. The large variance in sMSD has been found in the motion of chromatin loci in both *E.coli* and Human cells (49–53). To further quantify heterogeneities in the loci mobilities, we calculated the Van Hove function 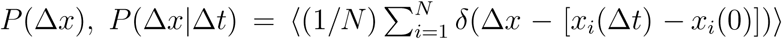. Figs.7C and 7D show the *P* (∆*x|*∆*t*) and normalized *P* (∆*x/σ|*∆*t*) for *ϵ* = 2.4*k*_B_*T* at different lag times ∆*t*. For *ϵ* = 1.0*k*_B_*T*, Van Hove function is well fit by a Gaussian at different lag times ∆*t* (Fig.S15). In contrast, for chromosome with glassy dynamics, all the *P* (∆*x|*∆*t*) exhibit fat tail, which is well fit by an exponential function at large values of ∆*x* (Fig.7C, D) at all *δt* values, suggestive of the existence of fast and slow loci (47).
5. The results in Fig.7 allow us to make direct comparisons with experimental data to establish signatures of dynamic heterogeneity. We calculated the distribution of effective diffusion exponent *α*_*i*_, *P* (*α*), where *α*_*i*_ is obtained by fitting the sMSD to ∼ *t^α^_i_* within some lag time (∆*t*) range. Fig.7E shows that *P* (*α*) calculated from simulations is in good agreement with experiments (54) in the same lag time range (0.42s < ∆*t* < 10s). The *P* (*α*) distribution in the range 10^−6^s < ∆*t* < 0.42s shows two prominent peaks, further validating the picture of coexisting fast and slow moving loci. The good agreement between the predictions of the CCM simulations with data, showing large variations of mobilities among individual loci *in vivo*, further supports our conclusion that organized chromosome dynamics is glassy. Interestingly, a recent computational study in which Human interphase chromosomes are modeled as a generalized Rouse chain suggests that the heterogeneity of the loci dynamics measured in live cell imaging is due to the large variation of cross-linking sites from cell to cell (24). Our model implies a different mechanism that the heterogeneity observed is a manifesto of the intrinsic glassy dynamics of chromosomes.

**FIG. 7:**
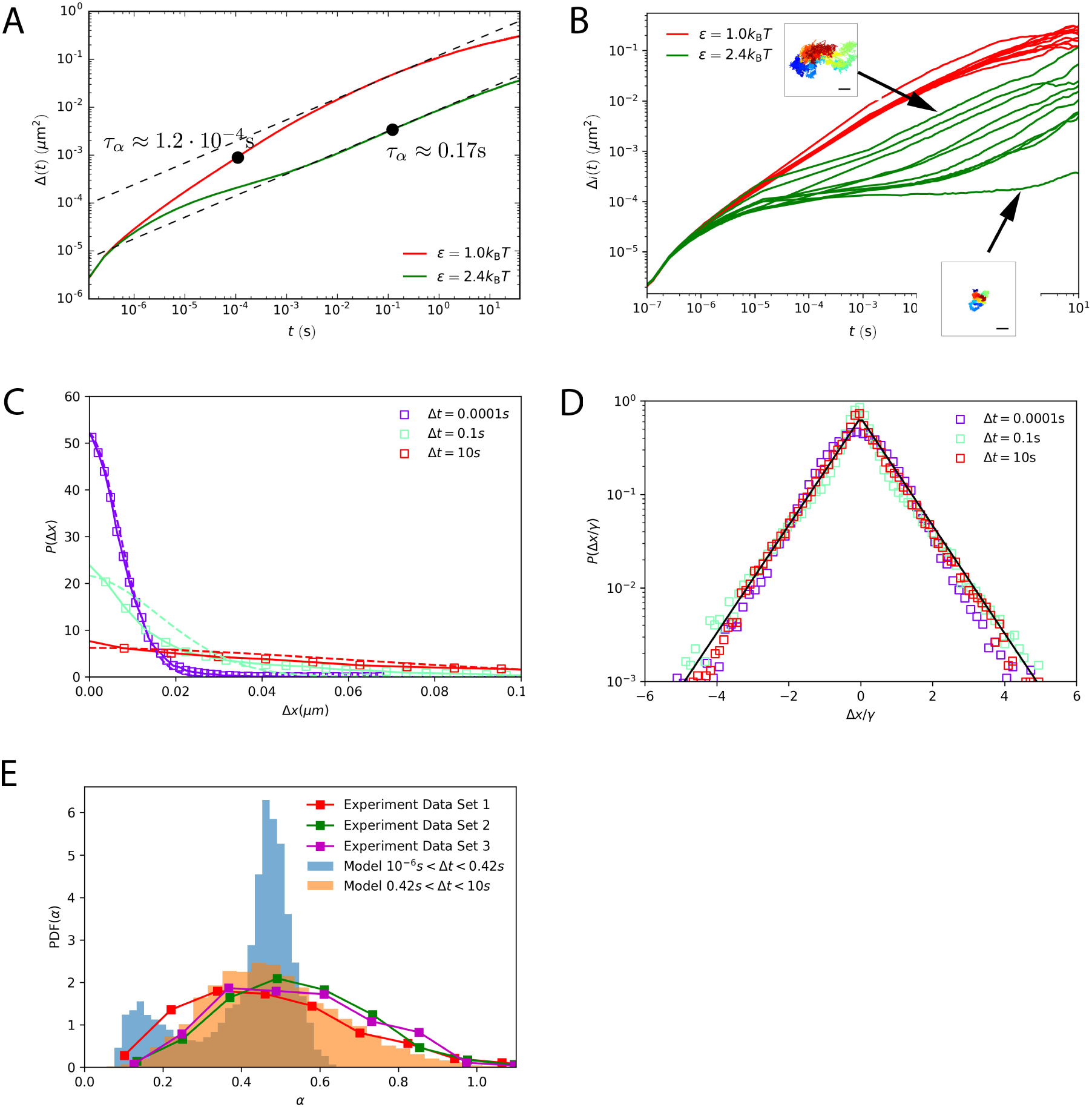
(top) **(A)** Mean Square Displacement, ∆(*t*), as a function of time, *t*. The effective diffusion coefficients, D, computed from the fitted dashed lines are 0.122*µ*m^2^/t^0.45^and 0.009*µ*m^2^/t^0.46^ for *ϵ* = 1.0*k*_B_*T* and *ϵ* = 2.4*k*_B_*T*, respectively. In the state exhibiting liquid-like behavior, the diffusion coefficient is more than one order magnitude larger than in the glassy state. Black dot marks the relaxation time *τ*_*α*_ defined as *F*_*s*_(*k, τ*_*α*_) = 1*/e* where *e ≈* 2.718. **(B)** Time dependence of 10 single loci MSD (sMSD, ∆_*i*_(*t*)) corresponding to 1^*st*^, 100^*th*^,…, 10, 000^*th*^ loci for *ϵ* = 1.0*k*_B_*T* and *ϵ* = 2.4*k*_B_*T*. The insets show ∆_*i*_(*t*) for two trajectories for fast (top) and slow (bottom) loci. Blue (red) indicates short (long) lag times. The scale bar is 1*µm*. Caging effect can be clearly observed as the plateau in ∆_*i*_(*t*). (**C**) The Van Hove function *P* (∆*x*) for *ϵ* = 2.4*k*_B_*T* at lag times ∆*t* = 0.0001s, ∆*t* = 0.1s and ∆*t* = 10s. *P* (∆*x*) has heavy tail at large ∆*x* and cannot be fit by a Gaussian (color dashed lines) except at ∆*t* = 0.0001s at small ∆*x*. **(D)** Same as **(C)** except displacement ∆*x* is normalized by the standard deviation *γ*. *P* (∆*x/γ*) for different lag times collapse onto a master curve. The black line is an exponential fit, ∼ *ϵ*^*−η*(∆*x/γ*)^ with *η ≈* 1.3.(**E**) Normalized distribution, *P* (*α*), of the effective diffusion exponent *α*. Comparison to experimental data (54) are also shown. The values of *α* are extracted from single loci trajectories by fitting sMSD, ∆_*i*_(*t*) ∼ *t*^*α*^. The lag time range 0.42s < ∆*t* < 10s is in the approximate same range used in the experiment. Experimental data set 1, 2, 3 are from Fig.2b, 2c, and Fig.S5 of Ref. (54), respectively. The results from our simulation (orange) agree well with experimental data, shown as orange. The blue bar plot is *P* (*α*) for small lag times 10^−6^s < ∆*t* < 0.42s. It shows two peaks, indicating the coexistence of two populations of loci with distinct mobilities.

### Active loci has higher mobility

Fig.8A shows MSD for active and repressive loci. For *ϵ* = 1.0*k*_B_*T*, there is no difference between active and repressive loci in their mobilities. However, in the glassy state active loci diffuses faster than the repressive loci. The ratio between the effective diffusion coefficients (the slope of the dashed line) of the active and repressive loci is 0.0116/0.008 *≃* 1.45, in good agreement with experimental estimate 0.018/0.013 *≃* 1.38 (32). Such a difference is surprising since the parameters characterizing the A-A and B-B interactions are identical. To investigate the origin of the differences between the dynamics of A and B loci, we plot the displacement vectors of the loci across the cross-section of the condensed chromosome (Fig.8B) for a time window ∆*t* = 0.1*s*. The loci on the periphery have much greater mobility compared to the ones in the interior. In sharp contrast, the fluid-like state exhibits no such difference in the mobilities of A and B (Fig.8D). To quantify the dependence of the mobility on the radial position of the loci, we computed the amplitude of the displacement normalized by its mean, as a function of the radial position of the loci, *r* (Fig.8C). For the chromosome exhibiting glass-like behavior, the mobility increases sharply around *r ≈* 0.7*µm* whereas it hardly changes over the entire range of r in the fluid-like system. Because the active loci are mostly localized on the periphery and the repressive loci are in the interior (Fig.2B), the results in Fig.8 suggest that the differences in the mobilities of the loci with different epigenetic states are due to their preferred locations in the chromosome. It is intriguing that glassy behavior is accompanied by a position-dependent mobility, which can be understood by noting that the loci in the interior are more caged by the neighbors, thus restricting their movement. In a fluid-like system, the cages are so short-lived that the apparent differences in the environments the loci experience are averaged out on short timescales. Note that in the experimental result (32) comparison is made between the loci in the periphery and interior of the nucleus. It is well known that the nucleus periphery is enriched with heterochromatin (repressive loci) and the interior is enriched with euchromatin (active loci).

**FIG. 8:**
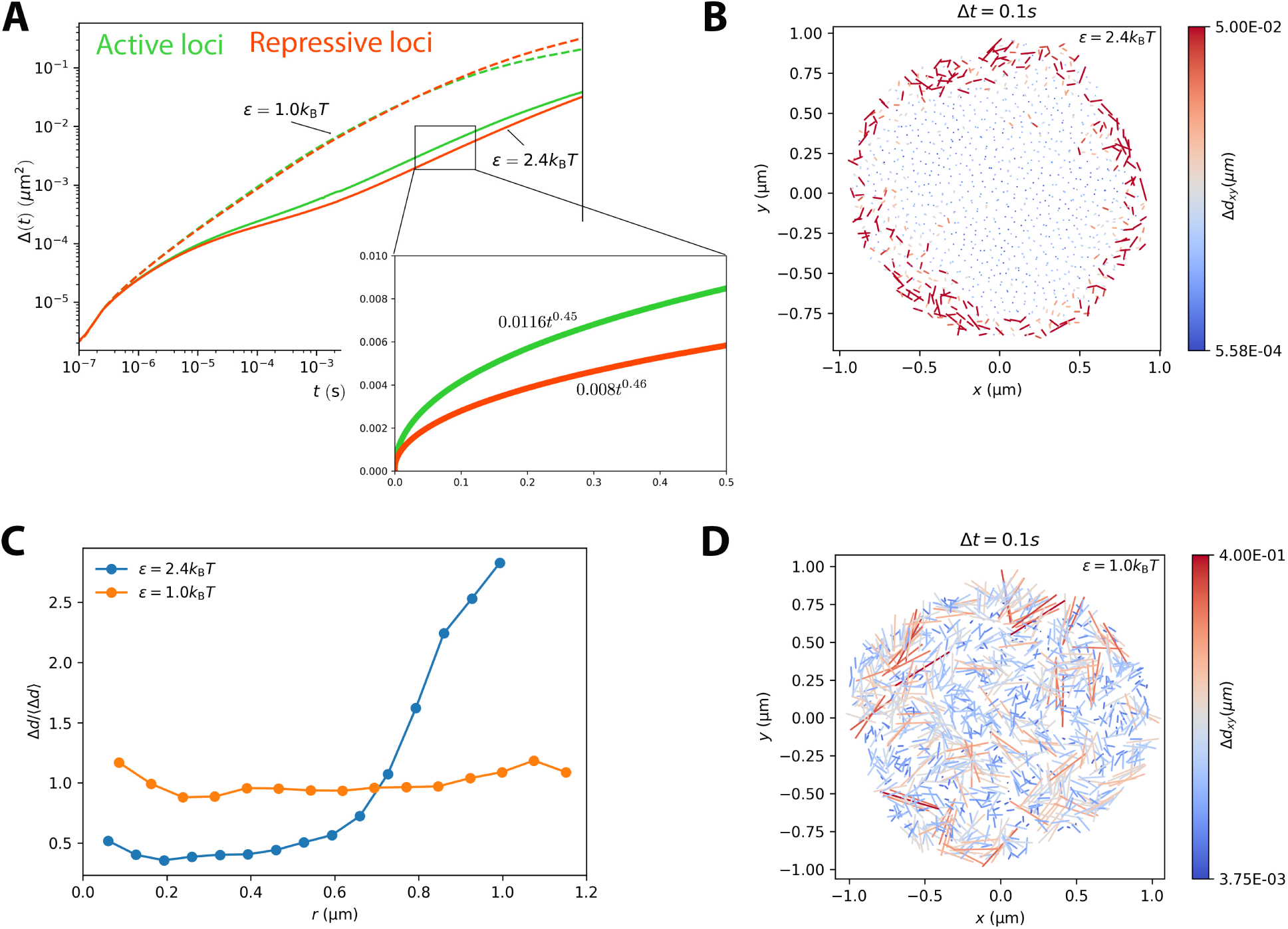
(A) The Mean Square Displacement for active loci and repressive loci. The equation shown in the inset is the fit using *Dt*^*α*^, where *D* is the diffusion coefficient and *α* is the diffusion exponent. **(B)** The displacement vectors of the loci within the equator cross-section of the structured chromosome for *ϵ* = 2.4*k*_B_*T*. The displacements are computed for time window ∆*t* = 0.1*s*. The color bars on the right show the magnitudes of the displacements. **(C)** Displacement ∆*d* normalized by its mean as a function of radial position, *r*, of the loci.**(D)** Same as (**B**) except the results are obtained using *ϵ* = 1.0*k*_B_*T*.

## Discussion

### Transferability of the CCM

In order to demonstrate the transferability of the CCM, we simulated Chr 10 using exactly the same parameters as for Chr 5. Fig.S17 compares the WLM obtained from simulations for different *ϵ* values and the computed WLM using the Hi-C contact map. The contact map is translated to the distance *R*_*ij*_ by assuming that 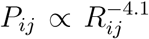 holds for Chr 10 as well. It is evident that the CCM nearly quantitatively reproduces the spatial organization of Chr 10 (Fig.S17). Thus, it appears that the CCM could be used for simulating the structures and dynamics of other chromosomes as well.

### Scale-dependent organization of chromosomes

In order to reveal how chromosome organizes itself and to link these processes to the experimentally measurable *P* (*s*), we calculated the time-dependent change in *P* (*s*) as a function of *t*. At scales less (above) than *s^∗^* ≈ 5*×*10^5^bps, *P* (*s*) decreases (increases) as the chromosome becomes compact. The *P* (*s*) ∼ *s*^−0.75^ scaling for *s < s*^*∗*^ (see also Fig.1B) is the result of organization on the small genomic scale during the early stage of chromosome condensation (Fig.9A). In the initial stages compaction starts by forming *≈ s*^*∗*^ sized chromosome droplets (CDs) as illustrated in Fig.9A. In the second scaling regime, *P* (*s*) ∼ *s*^−1.25^, global organization occurs by coalescence of the CDs (Fig.8A). Thus, our CCM model, which suggests a hierarchical chromosome organization on two distinct scales, also explains the two scaling in *P* (*s*).

**FIG. 9:**
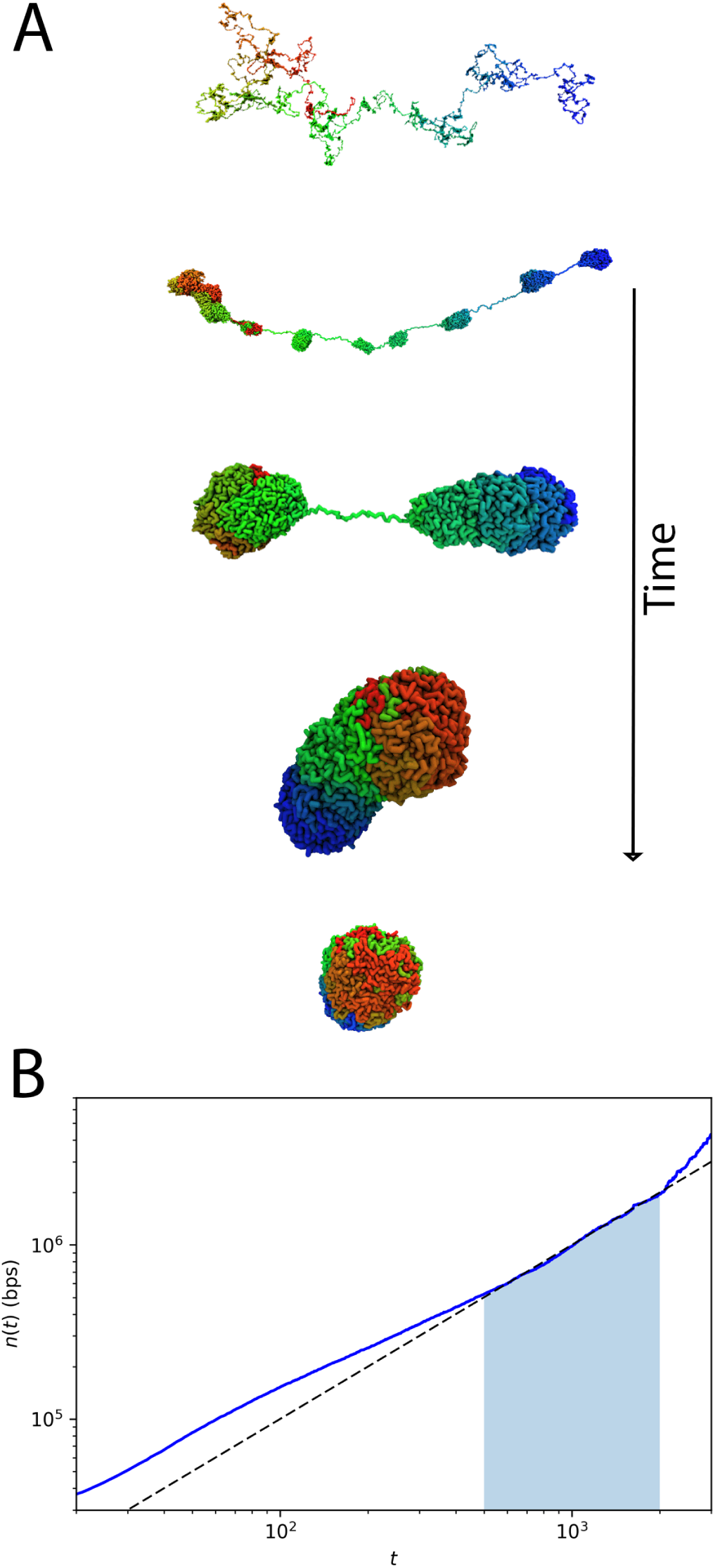
**(A)** Typical conformations sampled during the chromosome organization process, which begins by the formation of the chromosome droplets (CDs) connected by “tension strings”. Typical CD size, shown in the second subplot, is about *s ≈* 3 ⋅ 10^5^ ∼ 6 *⋅* 10^5^bps, which is consistent with the approximate value of *s*^*∗*^ (Fig.1B). CDs merge to form larger clusters. In the final stage, the two largest clusters condense to form the condensed chromosome. **(B)** The time-dependent growth of CDs, *n*(*t*), which is the average number of base pairs in a CD at *t*. The dashed line is a fit in the time window indicated by the shaded area, yielding *n*(*t*) ∼ *t*^1^. The roughly linear increase of *n*(*t*), over a range of times, is consistent with the Lifshitz-Slazov growth mechanism (55). For a vivid demonstration of this, see the movie in the SI.

The hierarchical nature of the structural organization is further illustrated using *A*(*s*), the number of contacts that a *s*-sized subchain forms with the rest of the chromosome. For a compact structure, *A*(*s*) ∼ *s*^2/3^ and *A*(*s*) ∼ *s* for an ideal chain. Fig.S18 shows that *A*(*s*) computed using the Hi-C data (black square line) varies as *s*^2/3^, suggesting that chromosome is compact on all length scales. We also find that upon increasing *ϵ*, the range of *A*(*s*) ∼ *s*^2/3^ expands. The pictorial view of chromosome organization (Fig.9A) and the *A*(*s*) scaling show that chromosome structuring occurs hierarchically with the formation of CDs and subsequent growth of the large CDs at the expense of smaller ones. We quantitively monitored the growth of CDs during the condensation process and found that the size of CD grows linearly with time during the intermediate stage (Fig.9B). Such a condensation process is reminiscent of the Lifshitz-Slazov mechanism (55) used to describe Ostwald ripening.

### Coincidence of scales

Our simulations show that the average TAD size and the crossover scale (*s*^*∗*^) the dependence of *P* (*s*) on *s* coincide. In addition, the size of the CDs is also on the order of *s*^*∗*^, which is nearly the same for all the chromosomes (Fig.S14). We believe that this is an important result. The coincidence of these scales suggests that both from the structural and dynamical perspective, chromosome organization takes place by formation of TADs, which subsequently arrange to form structures on larger length scales. Because gene regulation is likely controlled by the TADs, it makes sense that they are highly dynamic. We hasten to add that the casual connection between TAD size and *s*^*∗*^ as well as the CDs size has to be studied further. If this picture is correct then chromosome organization, at length scales exceeding about 100 Kbps, may be easy to describe.

## Conclusions

We developed the Chromosome Copolymer Model (CCM), a self-avoiding polymer with two epigenetic states and with fixed loop anchors whose locations are obtained from experiment to describe chromosome dynamics. The use of rigorous clustering techniques allowed us to demonstrate that the CCM nearly quantitively reproduces Hi-C contact maps, and the spatial organization gleaned from super-resolution imaging experiments. It should be borne in mind that contact maps are probabilistic matrices that are a low dimensional representation of the three-dimensional organization of genomes. Consequently, many distinct copolymer models are likely to reproduce the probability maps encoded in the Hi-C data. In other words, solving the inverse problem of going from contact maps to an energy function is not unique (see Ref. (9))

Chromosome dynamics is glassy implying that the free energy landscape has multiple equivalent minima. Consequently, it is likely that in genomes only the probability of realizing these minima is meaningful, which is the case in structural glasses. The presence of multiple minima also leads to cell-to-cell heterogeneity with each cell exploring different local minimum in the free energy landscape. We speculate that the glass-like landscape might also be beneficial in chromosome functions because only a certain minimum needs to be accessed to carry out a specific function, which will minimize large-scale structural fluctuations. In this sense, chromosome glassiness provides a balance between genomic conformational stability and mobility.

## Methods

### Construction of the CCM

Contact maps (4, 5) of interphase chromosomes show that they are partitioned into genome-wide compartments, displaying plaid (checkerboard) patterns. If two loci belong to the same compartment they have the higher probability to be in contact than if they are in different compartments. Although finer classifications are possible, compartments (4) can be categorized broadly into two (open (A) and closed (B)) classes associated with distinct histone markers. Open compartment is enriched with transcriptional activity-related histone markers, such as H3K36me3, whereas the closed compartment is enriched with repressive histone markers, such as H3K9me3. Chromatin segments with repressive histone markers have effective attractive interactions, which models HP1 protein-regulated interactions between heterochromatin regions (56, 57). We assume that chromatin fiber, with active histone markers, also has such a similar attraction. From these considerations, it follows that the minimal model for Human chromosome should be a copolymer where the two types of monomers represent open and closed chromatin states. To account for the two states, we introduce the Chromosome Copolymer Model (CCM) as a self-avoiding polymer with two kinds of beads. Models of the similar genre have been proposed in several recent studies (18–21, 27) to successfully decipher the organization of genomes.

The energy function in the CCM is, *E* = *U*_C_+*U*_LJ_, where *U*_C_ contains bond potential (*U*^S^) and loop interaction (*U*^L^), and *U*_LJ_ is the Lennard-Jones pairwise interaction between the monomers. If two monomers belong to type A (B), the interaction strength is *ϵ*_*AA*_(*ϵ*_*BB*_). The interaction strength between A and B is *ϵ*_*AB*_. Each monomer represents 1,200 base pairs (bps), with six nucleosomes connected by six linker DNA segments. The size of each monomer, *σ*, is estimated by considering two limiting cases. If we assume that nucleosomes are compact then the value of *σ* may be obtained by equating the volume of the monomer to 6*v* where *v* is the volume of a single nucleosome. This leads to *σ ≈* 6^1/3^*R*_N_ *≈* 20 nm where *R*_N_ *≈* 10 nm is the size of each nucleosome (58). Another limiting case may be considered by treating the six nucleosome array as a worm-like chain. The persistence length of the chromatin fiber is estimated to be ∼ 1,000 bps (59), which is about the size of one monomer. The mean end-to-end distance of a worm like chain whose persistence length is comparable to the contour length *L* is 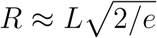. The value of *L* for a six nucleosome array is 6(16.5 + *R*_N_)nm where the length of a single linker DNA is 16.5nm. This gives us the upper bound of *σ* to be 130nm. Thus, the two limiting values of *σ* are 20 nm and 130 nm. We assume that the value of *σ* is an approximate mean, yielding *σ* = 70 nm.

The type of monomer is determined using the Broad ChromHMM track (60). There are totally 15 chromatin states, out of which the first eleven are related to gene activity. Thus, we consider state 1 to state 11 as a single active state (A) and states 12-15 as a single repressive state (B). For the genome range 146Mbps to 158Mbps in the Chromosome 5 in Human GM12878 cell, which is investigated mainly in this work, the numbers of active and repressive loci are 2369 and 7631, respectively. Details of the assignment are given in the Supplementary Information (SI).

### Simulations

As described in the SI, we performed simulations using both Langevin Dynamics (low friction) and Brownian Dynamics (high friction) using a custom modified version of the molecular dynamics package LAMMPS. The use of Langevin Dynamics accelerates the sampling of the conformational space (61), needed for reliable computation of static properties. Realistic value of the friction coefficient is used in Brownian Dynamics simulations to investigate chromosome dynamics, thus allowing us to make direct comparisons with experiments.

We varied the values of *ϵ*_*AA*_, *ϵ*_*BB*_ and *ϵ*_*AB*_ to investigate the effect of interaction strength on the simulation results. For simplicity, we set *ϵ*_*AA*_ = *E_BB_ ≡ E*. By fixing the ratio *E/E*_*AB*_ to 11/9, *ϵ* is the only relevant energy scale in the CCM. The results presented in the main text are obtained with *ϵ* = 2.4*k*_B_*T* unless stated otherwise. The contacts between loci in the simulation are determined by the threshold distance *r*_*c*_ = 2*σ* where *σ* = 70 nm

## Acknowledgements

We want to thank Abdul N Malmi-Kakkada and Xin Li for discussions and comments on the manuscript. We are grateful to the National Science Foundation (CHE 16-36424) and the Collie-Welch Regents Chair (F-0019) for supporting this work.

